# Functionally convergent but parametrically distinct solutions: Robust degeneracy in a population of computational models of early-birth rat CA1 pyramidal neurons

**DOI:** 10.64898/2026.03.30.715207

**Authors:** Matus Tomko, Carmen Alina Lupascu, Alzbeta Filipova, Peter Jedlicka, Lubica Lacinova, Michele Migliore

**Affiliations:** Centre of Biosciences, Institute of Molecular Physiology and Genetics, Slovak Academy of Sciences, Bratislava, Slovakia; Institute of Biophysics, National Research Council, Palermo, Italy; Biomedical Research Center, Cancer Research Institute, Slovak Academy of Sciences, Bratislava, Slovakia; Computer-Based Modelling in the Field of R Animal Protection, Faculty of Medicine, Justus Liebig University Giessen, Giessen, Germany

## Abstract

**Background:** Flexibility and robustness of neuronal function are closely linked to degeneracy, the ability of distinct structural or parametric configurations to produce similar functional outcomes. At the cellular level, this often manifests as ion-channel degeneracy, in which multiple combinations of intrinsic conductances yield comparable electrophysiological phenotypes.

**Methodology:** We used a population-based, data-driven modelling framework to generate large ensembles of biophysically detailed CA1 pyramidal neuron models constrained by somatic electrophysiological features extracted from patch-clamp recordings in acute slices from early-birth rats. 10 reconstructed morphologies were incorporated, and model populations were analyzed using parameter correlation analysis, principal component analysis, and generalization tests to assess robustness, degeneracy, and morphology dependence of intrinsic properties.

**Conclusions:** Across the model population, similar somatic firing behaviours emerged from widely different combinations of intrinsic parameters, demonstrating robust two-level ion channel degeneracy both within and across morphologies. Each morphology occupied a distinct region of parameter space, indicating morphology-specific compensatory effects, while weak pairwise parameter correlations suggested distributed compensation rather than tight parameter dependencies. Even with a fixed morphology, multiple parameter subspaces supported comparable electrophysiological phenotypes. Generalization across morphologies was structure-dependent and non-reciprocal, with successful parameter similarity occurring preferentially between structurally similar neurons. Interestingly, to accurately simulate spike-frequency adaptation, it was important to retain some kinetic properties of the ion channel models as free parameters during optimization. Together, these findings show that dendrite morphology shapes the valid parameter space, and similar electrophysiology of CA1 pyramidal neurons arises from the interplay between structural variability and ion-channel diversity. This work highlights the importance of population-based modelling for capturing biological variability and provides insights into how neuronal robustness might be maintained despite substantial heterogeneity, and offers a scalable pipeline for generating biophysically realistic CA1 neuron populations for use in network simulations.

**Author summary:** Neurons must reliably process information even though their internal components, such as ion channels and cellular shape, can vary widely from cell to cell. How stable behaviour emerges from such variability is a fundamental question in neuroscience. In this study, we explored this problem using detailed computer models of early-birth rat hippocampal CA1 pyramidal neurons, a cell type that plays a central role in learning and memory. Instead of building a single “average” neuron model, we created large populations of models that all reproduced key experimental recordings but differed in their internal parameters. We found that neurons with different shapes and different combinations of ion channels could nevertheless generate similar electrical activity. This phenomenon, known as ion channel degeneracy, allows neurons to remain functional despite biological variability or perturbations. Our results show that neuronal shape strongly influences which parameter combinations are viable, but that multiple solutions exist even for the same morphology. The population of models we provide offers a resource for future studies of early-birth CA1 pyramidal cell function and dysfunction.

## Introduction

Neurons express a wide variety of ion channels with diverse biophysical properties, providing a rich substrate for tuning intrinsic excitability. Even relatively small changes in ion-channel expression levels or spatial distributions can substantially alter neuronal activity [1]. Consistent with this sensitivity, large variability has been reported in electrophysiological measurements across cells of the same type, across experimental sessions, and across animals [1–5].

Degeneracy provides a key conceptual framework for understanding how reliable neuronal function can emerge from such variability. Degeneracy refers to the ability of structurally distinct elements or parameter combinations to yield similar functional outcomes, resulting in a many-to-one mapping between ion-channel configurations and neuronal or circuit phenotypes [6–8]. It is observed across multiple scales of brain organization and has been increasingly recognized as a fundamental principle underlying the flexibility and robustness of neural systems [6,9–11]. At the level of intrinsic physiology, ion-channel degeneracy enables different sets of coexpressed channels to play overlapping roles in shaping electrophysiological properties, thereby allowing compensation among conductances. This compensatory capacity renders neurons resilient to perturbations such as channel deletions, genetic mutations, expression noise, or environmental disturbances. In parallel, activity-dependent homeostatic mechanisms can further stabilize neuronal function by dynamically regulating ion-channel expression in response to deviations in physiological activity [2,12–14].

Biophysically detailed neuronal models allow systematic, reversible, and independent manipulation of morphological and biophysical parameters that are often inaccessible in experimental settings. Morphologically realistic models constrained by a broad set of physiological measurements are particularly important, as neuronal morphology imposes strong structural constraints on electrical signalling and synaptic integration [15–17].

Rather than adopting a one-size-fits-all approach in which a single model represents the average behaviour of a neuronal type [18], the population-based modelling framework constructs and validates large ensembles of biophysically realistic models that differ in their ion-channel configurations yet reproduce experimentally observed functional features [19–23]. This approach explicitly simulates biological variability and provides a framework for investigating degeneracy, parameter compensation, and robustness in neuronal systems. Population-of-models approaches have become an established tool for studying heterogeneity and differential parameter dependencies across multiple neuronal subtypes[6,17,20,24–27], including hippocampal CA1 pyramidal neurons [28].

CA1 pyramidal neurons occupy a critical position as the main output of the hippocampal circuit [29] and exhibit a rich and nonuniform repertoire of voltage- and calcium-dependent conductances distributed across their somatodendritic axis [28,30–32]. Despite this biophysical complexity, CA1 neurons reliably maintain characteristic firing patterns and integrative properties across animals and experimental conditions, suggesting the presence of robust compensatory mechanisms that stabilize intrinsic function.

Here, we simulate intrinsic electrophysiological properties using a unified, data-driven modelling workflow. Starting from experimental recordings and morphological reconstructions, we developed morphologically and biophysically detailed models of rat early-birth CA1 pyramidal neurons whose intrinsic properties are tightly constrained by experimental observations. Our results show that neuronal morphology strongly shapes the admissible parameter space, yet similar firing behaviours can emerge from distinct ion-channel configurations both across and within morphologies. Moreover, even for a fixed morphology, multiple parameter subspaces support comparable electrophysiological behaviour, demonstrating degeneracy in models of CA1 pyramidal neurons.

## Methods

### Experimental data

All experimental electrophysiological data used for this modelling study were obtained in our laboratory and have been previously published [33]. No new animal experiments were performed in the present study. Briefly, acute hippocampal slices were prepared from Wistar rats at postnatal days 11–13. Whole-cell patch-clamp recordings were performed in current-clamp configuration.

Single action potentials (AP) were activated using brief depolarizing current pulses (5 ms duration) with amplitudes increasing in steps of +50 pA. A series of APs was evoked by prolonged depolarizing current injections (300 ms duration) with amplitudes increasing in steps of +50 pA. Responses to hyperpolarizing current injection with a duration of 250 ms and amplitudes decreasing by −25 pA from a holding potential of - 70 mV were also recorded. Representative voltage responses and corresponding current injection for each stimulation protocol are shown in Figure 1a–c, and the electrophysiological features extracted across all current steps are listed in Supplementary Tables S1–S3.

**Figure 1:**
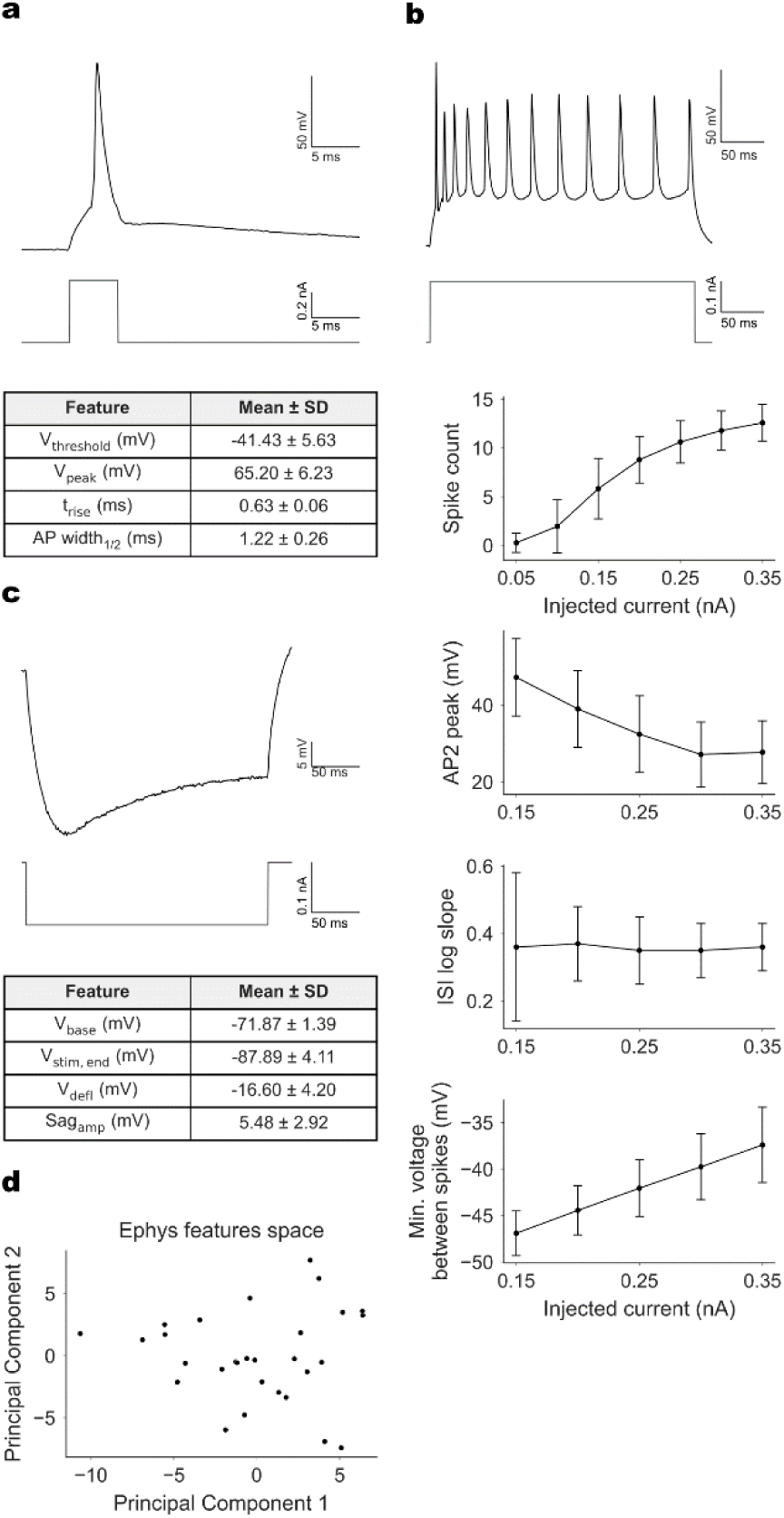
Experimental data used for the optimization pipeline. **(a)** Representative somatic voltage response to a brief depolarizing current pulse evoking a single action potential (top), corresponding stimulation protocol (middle), and selected electrophysiological features extracted across all recorded cells for this protocol (bottom). **(b)** Representative somatic voltage response to a prolonged depolarizing current pulse eliciting a train of action potentials (top), corresponding stimulation protocol (middle), and selected electrophysiological features extracted across all cells (bottom). **(c)** Representative somatic voltage response to a prolonged hyperpolarizing current pulse (top), corresponding stimulation protocol (middle), and selected electrophysiological features extracted across all recorded cells for this protocol (bottom). **(d)** Principal component analysis (PCA) of experimental feature vectors projected onto the first two components. Experimental cells form a single cluster.

Data from a total of 29 CA1 pyramidal neurons were included in the present study. Neurons were selected based on recording stability, absence of depolarization block, correct extraction of electrophysiological features, and availability of complete stimulation protocols required for model optimization. Full details of animal handling, slice preparation, recording conditions, and stimulation protocols are provided in [33].

Prior to feature extraction, all voltage traces were smoothed using a Savitzky–Golay filter to reduce high-frequency noise while preserving AP waveform characteristics. Filtering was performed using a custom-written Python script. Window lengths of 3 or 5 data points were applied around AP peaks, whereas a window length of 15 ms was used between APs or voltage traces without APs.

### Electrophysiological features

In principle, a large number of electrophysiological features (e-features) and model parameters can be considered during optimization. In practice, however, using all possible e-features is neither computationally feasible nor necessary. Therefore, we selected a subset of e-features that reliably capture the key characteristics of the experimental voltage responses (listed in S1–S3 Tables).

Extraction of e-features from somatic voltage traces was performed using the BluePyEfe Python package [34], which is an integral component of the BluePyEModel pipeline [35]. The selected e-features include descriptors of neuronal input–output behaviour, such as spike count and spike timing, as well as subthreshold features, including resting membrane potential, input resistance, and amplitude of voltage sag.

For each e-feature, the mean and standard deviation were computed across all 29 recorded neurons, yielding 182 e-features in total used to constrain the optimization process. A subset of e-features corresponding to current injection amplitudes not used during optimization was reserved for post-hoc generalization testing; these features are marked with an asterisk (*) in S1-S3 Tables.

Several global settings of BluePyEfe [34] were adjusted to match the experimental recording conditions. Voltage traces were interpolated with a temporal resolution of 0.1 ms. Spikes were detected when the membrane potential crossed 0 mV. For single AP responses, a derivative threshold of 15 mV/ms was used, and spikes occurring outside the stimulation window were included. For the AP series, a derivative threshold of 10 mV/ms was applied, and only spikes occurring during the stimulus period were considered.

### Updated stimulation protocols

Because the voltage recordings in the experimental data start 5 ms before the stimulus and end 30 ms after the stimulus, we updated the simulation protocols. The length of the stimuli was kept the same. However, we set delays of 100 ms for depolarizing pulses and 150 ms for hyperpolarizing pulses to ensure that all parameters stabilize. The simulations finished 45 ms after the stimulus for a brief 5-ms depolarizing protocol and 200 ms after prolonged depolarizing and hyperpolarizing protocols. In the case of the e-feature “steady_state_voltage” that represents the average voltage after the stimulus, the appropriate values of “voltage_base” (the average voltage during the last 10% of time before the stimulus) were used to keep a stable voltage after the stimulus.

### Morphologies

Because morphological reconstructions were not available for the neurons recorded in our laboratory, we used a set of 10 previously published morphologies of CA1 pyramidal neurons that matched the developmental age and rat strain of the experimental dataset [36]. All morphologies were obtained from the NeuroMorpho.org database. Only two of the selected reconstructions included an axon. To ensure consistency across morphologies, we therefore replaced axons in all models with a synthetic axon provided by BluePyOpt (bluepyopt.ephys.morphologies.NrnFileMorph) [37]. The synthetic axon consisted of two cylindrical sections, each 30 µm long and 1 µm in diameter. Axon replacement was implemented by the “bluepyopt_replace_axon” morphology modifier option in the recipes.json configuration file. Quantitative morphological characteristics of all reconstructions are summarized in S4 Table.

### Models configuration

All model parameters that were held constant across simulations are listed in S5 Table. Each model incorporated a set of ionic mechanisms characteristic of CA1 pyramidal neurons, as previously described in computational models [28,31]. The active membrane mechanisms included a transient sodium current (NaT), a persistent sodium current (NaP); four potassium currents (K_DR_, K_A_, K_M_, and K_D_); three voltage-gated calcium currents (CaN, CaL, and CaT); the hyperpolarization-activated nonspecific cation current (I_h_), and two calcium-dependent potassium currents (K_Ca_ and Cagk). Intracellular calcium dynamics were modelled using a simple extrusion mechanism with a single exponential decay time constant of 100 ms, applied to all compartments containing calcium channels. A complete list of mechanisms is provided in S6 Table.

Ion channels were distributed uniformly across dendritic compartments unless stated otherwise; consistent with experimental and modelling studies, the densities of K_A_ and HCN channels (mediating I_A_ and I_h_ currents) increased with distance from the soma [38,39]. The peak conductances of all ionic channels were treated as free parameters and independently optimized for each compartment type (soma, axon, basal dendrites, and apical dendrites). In addition, selected passive membrane parameters and ion channel kinetic parameters were optimized (S6 Table and params.json). Parameters bounds were chosen based on previously published models and refined by iterative trial-and-error to ensure stable firing behaviour and convergence of the evolutionary algorithm.

### Optimizations

Model optimizations were performed using BluePyOpt [37], an integral component of the BluePyEModel pipeline [35]. Parameter optimization employed the covariance matrix adaptation evolution strategy (CMA-ES), a stochastic optimization algorithm designed for non-linear, non-convex problems in continuous parameter spaces [40]. Offspring population sizes of 128 and 200 generations were used for all optimizations.

The optimization targeted a single objective, defined by a cost function known as the model’s total score. The total score was computed as the sum of individual e-feature Z-scores. For each e-feature, the Z-score was calculated as:

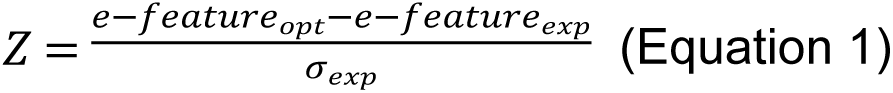

where 𝑒 ― 𝑓𝑒𝑎𝑡𝑢𝑟𝑒_𝑜𝑝𝑡_ and 𝑒 ― 𝑓𝑒𝑎𝑡𝑢𝑟𝑒_𝑒𝑥𝑝_ are the e-feature values from optimization and experiments, respectively, and 𝜎_𝑒𝑥𝑝_ is the standard deviation of the experimental e-features. All e-features contributed equally to the total score; no additional weighting was applied. Models were considered valid if all optimized e-features had Z-scores below 2.0. Thus, the experimental standard deviation serves as a feature-specific tolerance bound, ensuring that valid models remain within the range of biological variability observed across recorded neurons.

For each morphology, 10 independent optimization runs (seeds) were performed to generate distinct starting populations and explore alternative parameter solutions. Preliminary optimizations to refine parameter ranges and optimization settings were conducted on the CINECA Galileo100 high-performance computing infrastructure (Bologna, Italy). Final optimizations were executed on the DEVANA supercomputing system (Bratislava, Slovakia). Each optimization run used 60 CPU cores on a single node with 30 GB of memory and required ∼15 hours of wall-clock time, depending on the morphology.

### Generalization tests

We performed two generalization tests to assess model robustness. We use the term generalization rather than validation to distinguish these tests from approaches that evaluate models on entirely different protocol types; here, unseen current amplitudes within the same protocols are used to assess the model’s ability to generalize its responses.

First, a within-morphology generalization test was applied to all valid models obtained during optimization. In this test, all stimulation protocols were simulated, including current amplitudes used during optimization and additional current amplitudes not included in the fitting stage. Since the models are deterministic, e-features from current amplitudes used during optimization retain their original Z-scores below 2.0. Model validity for the generalization test was therefore assessed using only features extracted from the additional, unseen current amplitudes, and models were considered valid if all these features had absolute Z-scores below 3.0.

Second, a between-morphology generalization test was performed using the best-scoring model from each morphology. Optimized parameter sets were transferred to all remaining morphologies, all stimulation protocols were simulated, and total scores were computed. All simulations were executed on the DEVANA supercomputing system (Bratislava, Slovakia).

### Principal component analysis

Principal component analysis (PCA) was used to explore the structure and variability of models in both parameter space and electrophysiological feature space, as well as to characterize the experimental feature space. Prior to PCA, all datasets were standardized using Z-score normalization (StandardScaler, scikit-learn Python module).

PCA (PCA, scikit-learn Python module) was performed on all valid models pooled across morphologies. For parameter-space PCA, only optimized parameters were included. For model e-feature-space PCA, redundant e-features were removed prior to analysis by computing a pairwise correlation matrix; e-features with an absolute correlation coefficient greater than 0.9 were considered redundant. This procedure removed 82 e-features, resulting in a final set of 100 model e-features.

For PCA of experimental e-features, the proportion of missing values (NaNs) was first evaluated for each e-feature. E-features with more than 30% missing values were excluded (8 e-features). The remaining e-features were standardized using Z-score normalization, after which redundant e-features were removed using the same correlation-based criterion (|r| > 0.9), yielding a final set of 94 experimental e-features. The first two principal components were used for visualization and interpretation.

### Clustering analysis

Unsupervised clustering was performed using the K-means algorithm (KMeans, scikit-learn Python module) to identify groups of models with similar parameter or e-feature profiles, as well as the e-feature profiles of the experimental data. In the case of parameter space, clustering was performed in the reduced PCA space, where the number of principal components was set by the number that explained 85% of variance (18 components). For experimental and model e-features, clustering was performed on the same standardized datasets used for principal component analysis. The number of clusters (k) tested ranged from 2 to 10, and they were evaluated using the Silhouette score (silhouette_score, scikit-learn Python module) and Inertia. K-means was initialized multiple times with different random seeds to ensure robust clustering results. Cluster assignments were used for visualization and to assess whether models with distinct parameter combinations could produce similar electrophysiological behaviour, and experimental data are from a single phenotype.

### Statistical analysis

To compare differences in total score between models with different morphologies, a statistical test was performed. The Kruskal-Wallis test (kruskal, scipy Python module) was used, followed by Dunn’s post hoc test (posthoc_dunn, statsmodels Python module) with Holm correction. A value of p ≤ 0.05 was considered statistically significant.

### Ethics Statement

All experimental procedures were approved by the institutional ethical board and by the State Veterinary and Food Administration of the Slovak Republic, under permit No. 6604/2022, issued on July 26th, 2022. All procedures complied with Directive 2010/63/EU of the European Parliament and of the Council on the Protection of Animals Used for Scientific Purposes. Members of the institutional ethical board are Zuzana Bačová, Lucia Kršková, Katarína Danko, Gizela Gajdošíková, Ľubomír Vidlička, Martin Valachovič, and Jana Antalíková.

## Results

### Experimental data used for the optimization pipeline

To generate a population of computational models of early-birth CA1 pyramidal neurons, we employed the BluePyEModel pipeline [35], which consists of three main steps: (1) extraction of e-features, (2) parameter optimization, and (3) model generalization. In the first step, e-features were extracted from somatic voltage recordings (see Methods) obtained from 29 individual neurons. For each neuron, three stimulation protocols were applied: (1) brief 5 ms depolarizing current steps, (2) prolonged 300 ms depolarizing current steps, and (3) prolonged 250 ms hyperpolarizing current steps. In total, e-features were extracted from 4 voltage traces in the first protocol, 5 in the second, and 7 in the third, yielding 182 features per neuron. The mean values of the extracted e-features across all neurons were used as target values during optimization, while the corresponding standard deviations were used to normalize the e-feature–based scores. A summary of the average values and standard deviations for all e-features is provided in S1–S3 Tables.

Figure 1 shows representative voltage traces for each stimulation protocol, along with the average values of selected e-features. Figure 1a illustrates a typical AP elicited by a brief 5-ms depolarizing current pulse (0.5 nA) and the corresponding average values of selected spike-shape e-features. Figure 1b shows a series of APs characteristic of CA1 pyramidal neurons elicited by a prolonged depolarizing current pulse (0.2 nA). Consistent with classical electrophysiological characterizations of hippocampal pyramidal neurons [41], the number of spikes increased with injected current amplitude. In addition to increased spike count, neurons exhibited firing patterns characteristic of spike frequency adaptation during sustained depolarizing current steps. The peak amplitude of the second action potential decreased with increasing current amplitude, consistent with activity-dependent sodium channel inactivation reported in CA1 pyramidal neurons [42]. Despite higher firing rates, the logarithmic slope of the interspike interval remained stable across current amplitudes, indicating preserved adaptation dynamics. Moreover, the minimum membrane potential between spikes became progressively more depolarized with stronger input, consistent with elevated interspike voltages during strong depolarizing drive in CA1 neurons [43]. Figure 1c shows a typical membrane voltage response to hyperpolarized current injection (−0.125 nA) and average values of selected e-features, including a prominent voltage sag indicative of HCN channels activity typical of CA1 pyramidal neurons [36].

To assess whether the recorded neurons form a single electrophysiological phenotype, we performed PCA on the extracted e-features and then applied K-Means clustering. Based on the Silhouette score, the data were best described by a single cluster, indicating that all neurons belonged to a single phenotype. Figure 1d shows the projection of the e-features space onto the first two principal components.

### Optimization of CA1 pyramidal cell models

The second step of the BluePyEModel pipeline [35] is an optimization procedure that generates a population of models that reproduce the previously extracted e-features when simulated *in silico* under the same current stimulation protocols. The inputs to this step are: (1) detailed morphological reconstructions of neurons, (2) biophysical mechanisms describing passive and active membrane properties as well as intracellular ionic dynamics (S5-S6 Tables), and (3) experimentally derived e-features described in the previous section. Before optimization, stimulation protocols were divided into an optimization subset and a post hoc generalization test subset (S1-S3 Tables). Selected free parameters were optimized using an evolutionary algorithm (S5 Table). Model performance was evaluated using a cost function defined as the sum of Z-score–normalized errors between model-derived e-features and the corresponding experimental e-features’ mean values. The optimization procedure searched for parameter sets minimizing this cost function. Models for which all e-features used for optimization had absolute Z-scores below 2.0 were classified as valid. These models were subsequently subjected to the post hoc generalization test and included in downstream analyses.

For the optimization, we used 10 detailed morphological reconstructions of CA1 pyramidal neurons from Marcelin et al. (2012) [36], obtained from the NeuroMorpho.org database. The selected morphologies matched the developmental age range and Wistar rat strain of the experimental recordings used to extract the target e-features. Quantitative morphological characteristics are summarized in the S4 Table. Figure 2a shows the morphologies used in this study (top row) along with their identifiers, as well as example voltage traces from the best-scoring model for each morphology in response to a representative current step from each of the three stimulation protocols. APs elicited by brief depolarizing current pulses exhibited similar waveforms across morphologies and closely matched the experimentally recorded spike shape. During prolonged depolarizing current injection, the optimized models produced a series of APs with qualitatively similar firing patterns across morphologies and experimental data. In particular, the amplitude of the second AP was consistently lower than that of the first AP, consistent with activity-dependent sodium channel inactivation reported in CA1 pyramidal neurons [42]. Additionally, the minimum voltage between spikes increased with increasing current amplitude, reproducing a feature observed in the experimental data, reported for CA1 pyramidal neurons but often not explicitly constrained in computational models [43]. In response to hyperpolarizing current injection, all models exhibited a pronounced voltage sag indicative of HCN channel activation. Together, these results demonstrate that the evolutionary algorithm successfully identified morphology-specific parameter sets that generate consistent and biologically realistic voltage responses across diverse CA1 pyramidal neuron morphologies.

**Figure 2.**
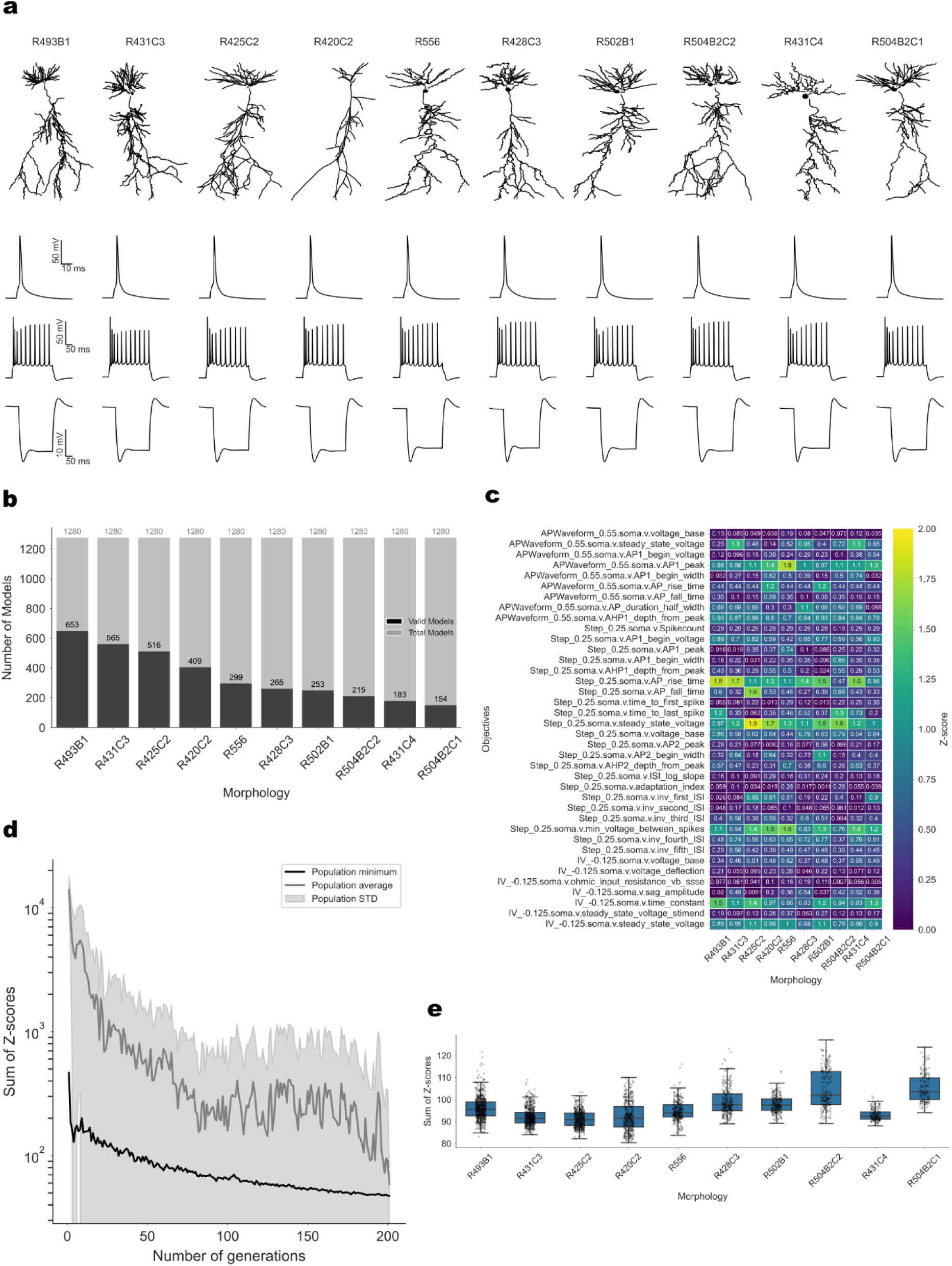
Optimization of CA1 pyramidal cell models. **(a)** Three-dimensional reconstructions of the ten CA1 pyramidal cell morphologies used in this study (from Marcelin et al., 2012) and representative somatic voltage traces from the best-scoring model of each morphology across three stimulation protocols: brief depolarizing pulse (0.55 nA, 5 ms; top), prolonged depolarizing pulse (0.25 nA, 300 ms; middle), and prolonged hyperpolarizing pulse (−0.125 nA, 250 ms; bottom). **(b)** Number of valid models obtained in the final generation for each morphology (10 seeds per morphology, offspring size = 128; validation threshold = 2 for features used in optimization and 3 for features contributing to validation/generalization). **(c)** Heatmap of Z-scores for selected optimization features for the best-scoring model of each morphology. **(d)** Example of the evolution of the total score during a single optimization run (corresponding to the run that produced the overall best model). **(e)** Boxplots of total Z-scores (sum across all features), with individual model scores overlaid as points. Some morphologies systematically achieve lower (better) scores, whereas others perform less well; several morphology pairs exhibit significant differences in total score distributions.

Using this approach, we generated a large population of valid CA1 pyramidal neuron models. A model was considered valid if all e-features used during optimization had Z-scores below 2 and all e-features evaluated during the post-hoc generalization test (unseen during optimization) had Z-scores below 3.

Figure 2b shows the total number of valid models grouped by morphology. The largest number of valid models was obtained for the “R493B1” morphology (653 models), with a gradual decrease across morphologies, with “R4504B2C1” morphology yielding 154 valid models. Spearman correlation analysis between morphological properties (S4 Table) and the number of valid models (Figure 2b) revealed moderate positive correlations with measures of dendritic complexity, including total length (ρ = 0.49), number of sections, branches, and bifurcations (ρ = 0.47 each), number of segments (ρ = 0.39), and maximal path distance (ρ = 0.38). A moderate negative correlation was observed with fractal dimension (ρ = -0.40), and negligible correlations with remaining properties (|ρ| < 0.27). However, none of these correlations reached statistical significance (all p > 0.05), likely reflecting limited statistical power given the small number of morphologies (n = 10).

Figure 2c shows the Z-scores of the best-scoring model for each morphology corresponding to the voltage traces shown in Figure 2a. For responses to brief depolarizing current steps, all models showed low Z-scores for most e-features, indicating robust reproduction of single-spike properties. During prolonged depolarizing current injection, all models except the “R504B2C2” morphology reproduced the experimentally observed number of APs. Across morphologies, most e-features were well optimized, whereas higher Z-scores were consistently observed for AP rise time, steady-state voltage, and the minimum voltage between spikes. In response to hyperpolarizing current injection, all models exhibited a clear voltage sag, with elevated Z-scores primarily associated with the membrane time constant.

Figure 2d shows an example of the evolution of the total score as a function of the number of generations during the optimization process. The figure illustrates an optimization run from which the best-performing model was obtained. The optimization gradually converged to a stable solution, with a marked decrease in the population-average score around generation 180. Extending the optimization to 200 generations enabled further gradual improvement of the best-performing models, resulting in lower final total scores. Figure 2e shows the distribution of total Z-scores grouped by morphology. The “R425C2” and “R431C3” morphologies exhibited the lowest median total scores, while “R504B2C1” and “R504B2C2” showed the highest median scores and largest interquartile range. Notably, a low total score did not necessarily correspond to a higher number of models, as validity depended on individual feature Z-scores rather than the total score alone. Significant differences in total score between morphologies were detected using the Kruskal–Wallis test (H=1251.97, p=7.293×10^-264^), and pairwise post hoc comparisons were performed using Dunn’s post hoc test with Holm correction for multiple comparisons (S7 Table).

Taken together, these results demonstrate that the optimization procedure generated a large population of valid CA1 pyramidal neuron models that require morphology-specific parameter solutions (see the next sections). To further assess whether these optimized models reproduce key functional properties observed experimentally, we next examined their input-output relationships.

An informative summary of how the optimized models capture experimental input–output properties is provided by the input/output (I/O) relationship. Figure 3a shows the number of action potentials as a function of somatic current injection amplitude for experimental data (mean ± SD) and for the best-performing model of each morphology. Across morphologies, the model I/O curves closely overlapped with the experimental data for current steps of 0.05 nA and for currents ≥ 0.2 nA, including stimulation amplitudes used exclusively for the post-hoc generalization test. Minor deviations were observed near threshold, where spike initiation is particularly sensitive to intrinsic variability. Moreover, all best-performing models exhibited comparable total Z-scores, ranging from 80.39 (“R420C2”) to 94.03 (“R504B2C1”). Together, these results indicate that the optimized models robustly reproduce the experimentally observed input/output properties across morphologies.

**Figure 3.**
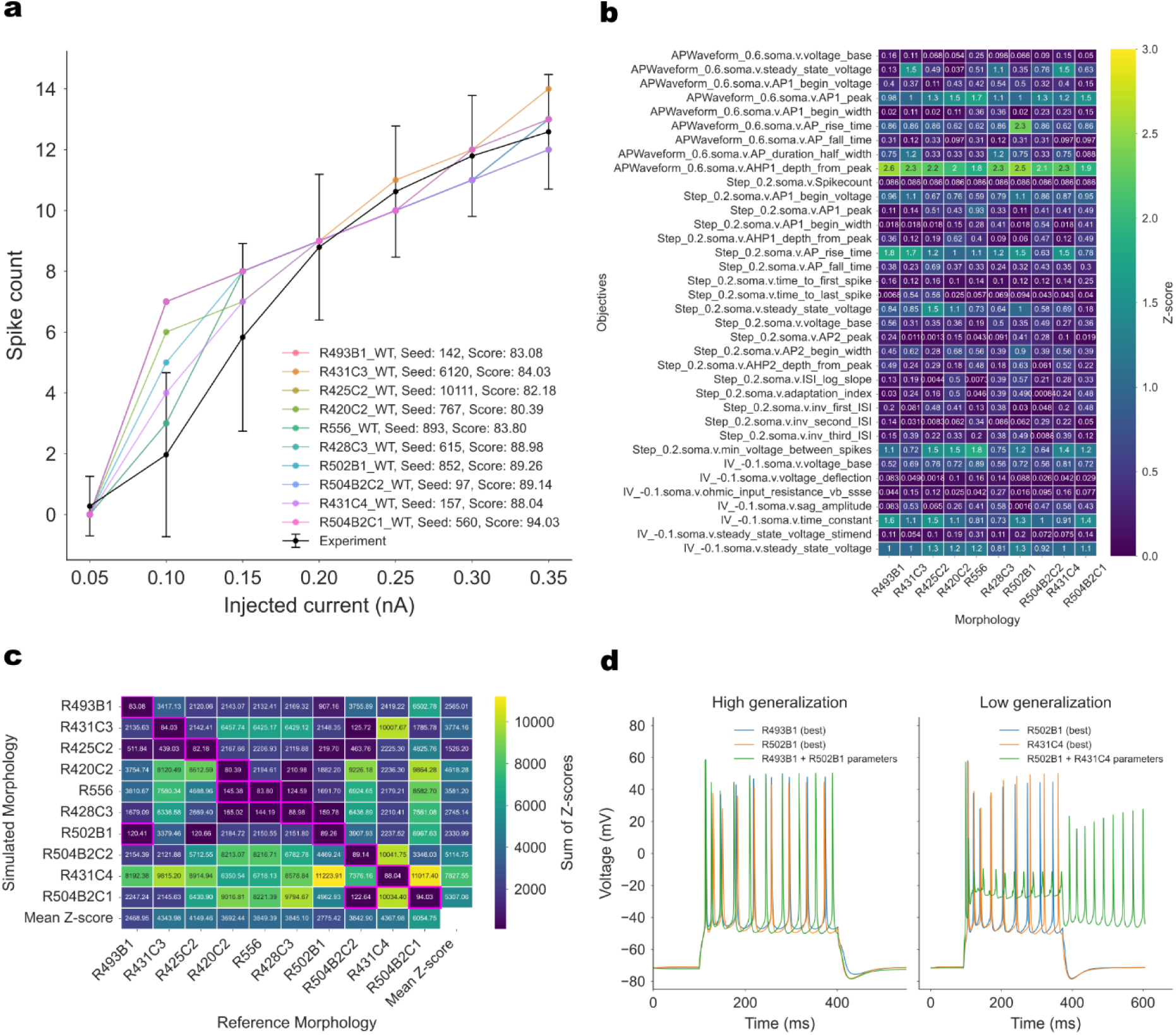
Generalization tests of optimized CA1 pyramidal cell models. **(a)** Input–output (I/O) relationships of the best-scoring model from each morphology (colored traces) compared with experimental data (black; mean ± standard deviation). **(b)** Heatmap of scores for selected features used during the generalization test, computed for the best-scoring model of each morphology. **(c)** Cross-morphology generalization matrix. For each best-scoring model, its optimized parameter set was applied to all remaining morphologies, and the resulting feature scores are shown. Magenta rectangles indicate cases in which the model passed validation. The final column reports the mean score across simulated morphologies (reflecting how well each morphology receives parameters), whereas the final row reports the mean score obtained when each morphology provides parameters. Overall, R425C2 shows the best generalization across simulated morphologies, while R493B1 provides parameters that generalize most effectively. **(d)** Representative somatic voltage traces illustrating cases of high and low cross-morphology generalization.

### Generalization tests of optimized CA1 pyramidal cell models

To assess the robustness of our optimization approach, we conducted two types of generalization tests: (1) within-morphology and (2) cross-morphology. The within-morphology generalization test is an integral part of the BluePyEModel pipeline [35] and was implemented by selecting subsets of current injection amplitudes from each stimulation protocol that were excluded from the optimization and evaluated only in a post-hoc test. Figure 3b shows a heatmap of Z-scores for selected e-features obtained from one representative current step per protocol used in the generalization test, shown for the best-performing model of each morphology. For the majority of e-features, Z-scores remained below 1.0 across morphologies, indicating good generalization to unseen stimulation amplitudes. The only feature that consistently exhibited elevated Z-scores (>2.0) was the afterhyperpolarization depth from peak. Overall, these results demonstrate that the optimized models generalize well to current steps not used during optimization.

To evaluate cross-morphology generalization, we transferred the optimized parameter sets from the best-performing model of each morphology to all remaining morphologies and evaluated their performance using the same validation criteria. Figure 3c summarizes this analysis by total Z-score, with magenta rectangles indicating models that passed validation. In addition to the diagonal elements corresponding to within-morphology optimization, three off-diagonal cases exhibited successful cross-morphology generalization in one direction but not in the reverse direction. To quantify generalization performance, we computed the mean total Z-score for each morphology when used as a target morphology (column-wise mean) and when its parameters were applied to other morphologies (row-wise mean). Among the tested morphologies, “R493B1” exhibited the lowest column-wise mean Z-score (2468.95), indicating the highest suitability as a reference morphology, whereas parameter sets optimized for “R425C2” produced the lowest row-wise mean Z-score (1526.20), indicating superior generalization across morphologies.

To further illustrate the range of generalization outcomes, Figure 3d shows example voltage traces for cases of high and low cross-morphology generalization. In the high-generalization case, applying parameters optimized for the “R502B1” morphology to the “R493B1” morphology yielded nearly identical voltage responses, whereas the reverse transfer did not produce a valid model. In contrast, in the low-generalization case, applying parameters optimized for the “R431C4” morphology to the “R502B1” morphology led to depolarization block during current injection and sustained firing following stimulus offset. Taken together, these results demonstrate robust within-morphology generalization and reveal that morphological differences strongly constrain the transferability of optimized parameter sets across CA1 pyramidal neuron morphologies.

### Degeneracy in the CA1 pyramidal neurons models

Degeneracy refers to the phenomenon whereby distinct combinations of model parameters can reproduce similar experimentally observed firing properties across stimulation conditions, a property that has been widely reported in both biological neurons and conductance-based models [6,8,20,28]. To assess degeneracy in our model population, we analysed both the parameter space and the corresponding e-feature space.

Figure 4a shows violin plots of normalized parameter distributions across all valid models, with coloured dots indicating parameter values of the best-performing model for each morphology. Many parameters exhibited broad distributions spanning most of their allowed ranges, with best-performing models distributed across these ranges, indicating substantial parameter degeneracy and morphology-dependent shaping of parameter space. In contrast, several parameters showed narrower distributions or were consistently constrained near parameter bounds or central values, including the maximal conductance of the HCN channel in all compartments except axon (ghdbar.hd_allnoaxon), maximal conductance of the transient somatic sodium channel (gbar.nax_somatic), and the axonal proximal A-type potassium channel (gkabar.kap_axonal),, with best models clustering tightly. These parameters are therefore likely to play a more critical role in reproducing the experimental firing properties.

**Figure 4.**
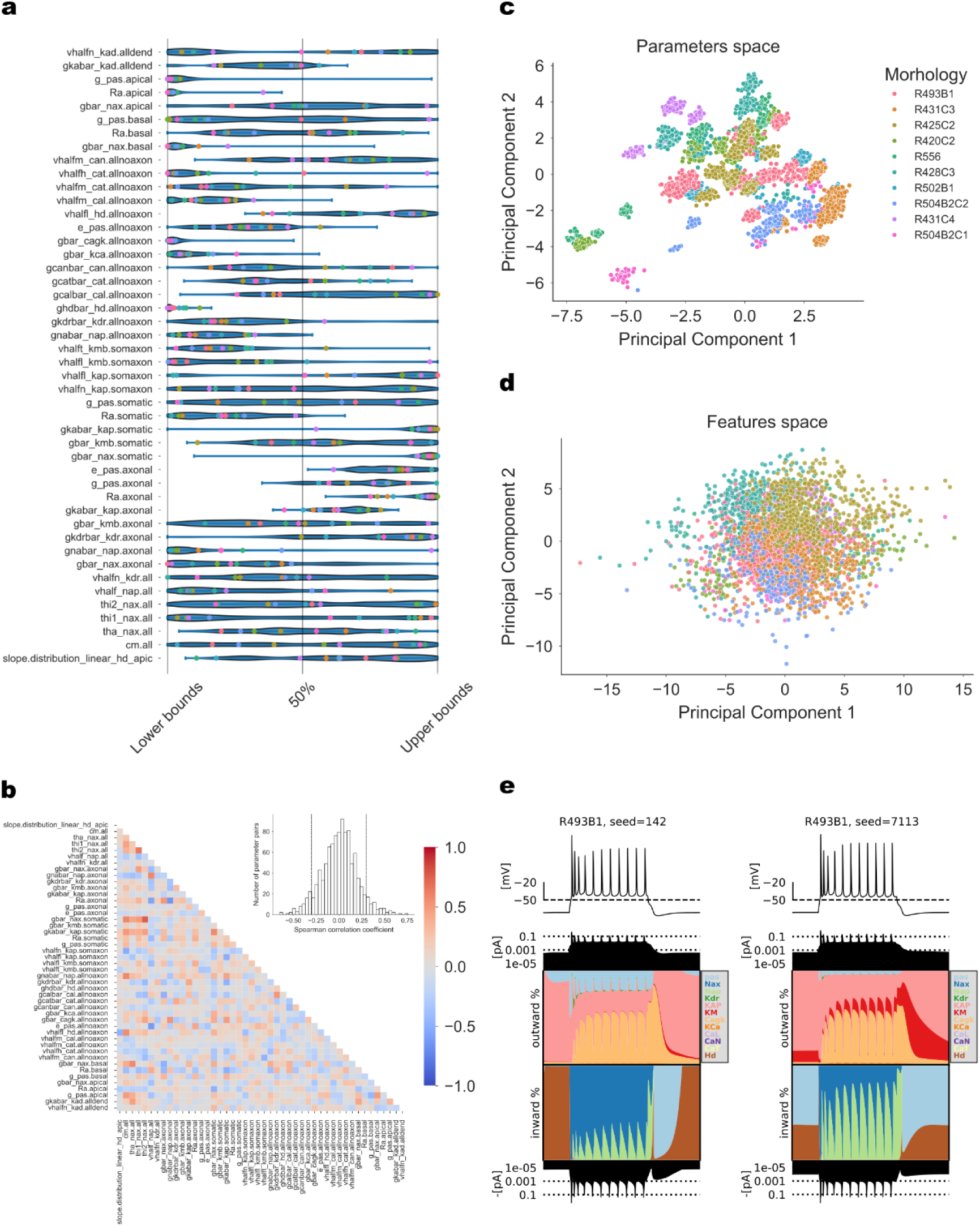
Degeneracy in CA1 pyramidal neuron models. **(a)** Normalized optimized parameter values for all valid models. Colored dots indicate parameter values of the best-scoring model for each morphology. **(b)** Spearman correlation heatmap of all parameters, revealing weak correlations among multiple parameter pairs, consistent with parametric degeneracy. Histogram of correlation coeficients in upper right corner. **(c)** Principal component analysis (PCA) of parameter vectors projected onto the first two components. PCA reveals morphology-dependent structure in parameter space, with distinct subregions and subclusters within individual morphologies. **(d)** PCA of feature vectors (82 redundant features removed; 100 retained) projected onto the first two components. Feature space analysis shows convergence of all models toward a single electrophysiological phenotype across morphologies, consistent with experimental observations and the principles of degeneracy. KMeans clustering yielded a single cluster based on the Silhouette score. **(e)** Currentscape comparison for two optimized R493B1 models originating from different optimization seeds and parameter-space PCA subclusters, but exhibiting similar total scores and voltage responses. Despite nearly identical firing patterns, the relative contributions of ionic currents differ, illustrating biophysical degeneracy within a single morphology.

To further assess compensatory relationships among parameters, we computed pairwise Spearman correlation coefficients for all 1035 unique parameter pairs, summarized as a correlation heatmap in Figure 4b, together with the distribution of correlation coefficients. Strong positive or negative correlations indicate potential compensatory interactions, whereas weak correlations indicate parametric independence and degeneracy. The distribution of correlation coefficients revealed that most parameter pairs were weakly correlated (|ρ| < 0.3 for 87% of all pairs, 901/1035), consistent with a highly degenerate parameter space. Only a small subset of parameter pairs exhibited strong correlations, suggesting that model validity does not rely on tight one-to-one parameter dependencies. Notably, membrane capacitance showed strong correlations with multiple parameters, consistent with its central role in shaping membrane dynamics in morphologically detailed models. In addition, peak sodium conductance was correlated with sodium channel kinetic parameters, reflecting known trade-offs in sodium channel availability and excitability.

To better characterize the structure in parameter and feature spaces, we performed PCA. PCA of normalized parameter vectors revealed that the first 16 principal components accounted for approximately 90% of the total variance. K-means clustering of these components did not identify a clear multi-cluster structure (Silhouette score < 0.5 for all tested cluster numbers), suggesting an overall continuous parameter manifold. Nevertheless, inspection of the first two principal components showed that models corresponding to different morphologies occupied partially distinct but overlapping subspaces, and that within each morphology, models formed multiple subclusters. This indicates that the optimization process identified multiple alternative parameter solutions for a given morphology. In contrast, PCA performed on the e-feature space revealed a single compact cluster (Figure 1d), consistent with the single-cluster structure observed in the experimental data.

To further illustrate the biophysical basis of degeneracy within a single morphology, we compared ionic current contributions for two “R493B1” models with similar total scores (83.08 and 84.97) but originating from different optimization seeds and occupying distinct subclusters in parameter-space PCA. Despite their different parameter configurations, both models exhibited highly similar voltage responses to the same current injection (Figure 4e). However, the underlying ionic current contributions differed substantially, with distinct relative contributions of sodium, potassium, calcium, and Ih currents during spiking and interspike intervals. This demonstrates that multiple, mechanistically distinct parameter combinations can generate equivalent electrophysiological behaviour within the same morphology, providing a biophysical explanation for the observed subclustering in parameter space. Together, these results demonstrate two-level degeneracy in the model population: distinct morphologies and multiple parameter configurations within each morphology can give rise to similar electrophysiological behaviour.

## Discussion

In this work, we developed a population of conductance-based and morphologically detailed models of early-birth rat CA1 pyramidal neurons that reproduce key electrophysiological responses observed in patch-clamp experiments. Using the BluePyEModel pipeline [35] models were optimized to match experimental firing properties, including the shape of single action potential, repetitive firing, and responses to hyperpolarizing current injection. The models were subsequently evaluated using within- and cross-morphology generalization tests. By combining multiple detailed neuronal morphologies with independent optimization runs, we generated a large population of valid models that accurately reproduced mean experimental electrophysiological features while satisfying stringent feature-based constraints, though the variability across the model population was lower than observed experimentally (see limitations).. Our results demonstrate that neuronal morphology strongly shapes the admissible parameter space, yet similar firing behaviours can emerge from distinct parameter combinations both across and within morphologies. PCA further revealed that, even for a fixed morphology, multiple parameter subspaces can support comparable electrophysiological behaviour, providing clear evidence for robust two-level degeneracy in early-birth rat CA1 pyramidal neuron models.

The optimized model population faithfully reproduces several hallmark somatic electrophysiological properties of early-birth CA1 pyramidal neurons observed in our experimental recordings [33]. These include realistic single-AP waveforms, spike-frequency adaptation during sustained depolarization, and prominent voltage-sag responses to hyperpolarizing current injections, consistent with established experimental characterizations of CA1 pyramidal cells [28,41]. Importantly, models captured not only spike counts across input amplitudes but also subtler features, such as activity-dependent reduction of action potential peak amplitude, stable interspike interval adaptation slopes, and progressive depolarization of interspike minima under stronger depolarizing drive, all of which are well documented in hippocampal pyramidal neurons [36,42,43].

In our optimization framework, all electrophysiological features were weighted equally, such that no single aspect of the response was prioritized a priori. This choice reflects an intentional emphasis on general reproduction of neuronal behaviour rather than preferential fitting of specific features, such as spike count or AP shape. Therefore, the resulting models achieve good overall agreement with experimental data across multiple stimulation protocols, while tolerating moderate deviations in individual features. Moreover, allowing selected kinetic parameters to vary during optimization proved essential for capturing experimentally observed spike timing and adaptation dynamics, particularly under sustained stimulation. Together, these results demonstrate that accurate reproduction of CA1 pyramidal neuron input–output behaviour can be achieved through multiple parameter combinations, without overfitting individual features or protocols.

A main outcome of our work is the extensive degeneracy observed across optimized CA1 pyramidal neuron models, whereby similar somatic electrophysiological phenotypes emerged from widely different combinations of intrinsic parameters. Such degeneracy has been repeatedly reported across neuronal types and levels of analysis and is now recognized as a fundamental organizing principle of neuronal function rather than a modelling artifact [2,6,20,24–26,44–46]. Experimental studies have demonstrated that neurons exhibit highly variable levels of ion channels and related genes while maintaining functional outputs [5,46], and computational studies have shown that this variability can be reconciled with the maintained outputs through compensatory interactions among conductance and kinetics [14,23,26–28,47]. Consistent with these findings, our analysis revealed that most pairs of optimized parameters were only weakly correlated, indicating that model validity does not depend on tightly constrained one-to-one parameter relationships but instead arises from many-to-one mappings between parameter space and electrophysiological behaviour [14,48].

Importantly, our results demonstrate that degeneracy is strongly modulated by neuronal morphology. While valid models from different morphologies occupied overlapping regions of e-feature space (Figure 4d), their corresponding parameter distribution and PCA cluster structures were morphology-specific, indicating that morphology shapes the accessible regions of intrinsic parameter space (Figure 4a, c). This finding aligns with recent modelling studies showing that morphology and biophysics jointly constrain neuronal excitability, rather than acting as independent factors [17,27,28,49,50]. Previous works in CA1 pyramidal neurons have reported substantial variability in channel densities across morphologically distinct cells while preserving functional output [28,51,52]. Our results extend these observations by systematically quantifying how multiple morphologies give rise to distinct but degenerate parameter ensembles using the same mean experimental neuron. Thus, morphology acts both as a source of variability and as a structural constraint that shapes the landscape of compensatory solutions available to a neuron.

In addition to degeneracy across morphologies, our analyses reveal pronounced degeneracy within individual morphologies. PCA and clustering of optimized parameters showed that, even for a fixed dendritic structure, multiple distinct parameter clusters emerge that yield comparable model performance (Figure 4c). Using currentscape plots (Figure 4e) [53], we demonstrate that models with nearly identical electrophysiological scores can rely on different ionic current compositions, reflecting potential alternative compensatory strategies at the level of individual channels. Similar findings have been reported in other detailed neuron models, where accurate functional output was maintained despite substantial rebalancing of ionic currents or even removal of specific conductances [23,27,28]. Together, these results highlight that model validity does not depend on a unique ionic mechanism, but rather on a flexible interplay among multiple conductances within morphology-specific constraints.

Biological neurons must remain functional despite ongoing perturbations, including variability in channel expression, and cellular structure, while retaining sufficient flexibility to operate across a broad dynamic range [7,25,45]. To assess robustness and generalization of the optimized models, we performed two complementary generalization tests.

The first test followed a train-test paradigm commonly used in machine learning, in which models are evaluated on stimuli not encountered during optimization. Specifically, a subset of current injections from each protocol was excluded from the optimization phase and used exclusively for generalization, with model validity assessed using a Z-score threshold of 3 for all extracted e-features. The majority of valid models successfully passed this test, indicating that the optimization did not overfit the training stimuli but captured intrinsic electrophysiological properties that generalize to unseen inputs.

In the second test, we assessed cross-morphology generalization by transferring parameter sets from the best-performing models of one morphology to other morphologies. In contrast to some previous studies that reported broader parameter transferability [27,28], but in line with previous modelling of hippocampal granule cells [49] generalization across morphologies in our dataset was limited and not reciprocal.

This outcome is not unexpected, given (1) the large number of free parameters optimized, (2) the focus on best-performing models rather, and (3) the observation that high-performing models may belong to distinct, non-overlapping clusters in parameter space. Together, these results highlight morphology as a strong structural constraint that shapes the set of admissible compensatory solutions, rather than a neutral background on which parameters can be freely transferred.

Several limitations of the present study should be acknowledged. First, the models focus exclusively on intrinsic electrophysiological properties under somatic current injection and do not incorporate synaptic inputs or dendritic recordings. Second, the morphologies used in this study were obtained from previously published datasets and were not reconstructed from the same neurons used for electrophysiological recordings, although they were selected to match the rat strain and developmental age of the experimental animals.

Third, despite generating a large population of optimized models, the full extent of experimental variability may not be completely captured. This limitation arises from the feature-based optimization framework and the way it handles missing features. For example, at a depolarizing current step of 0.1 nA, some experimental neurons fired action potentials while others did not. Because the mean experimental spike count for this stimulus was 1.96 ± 2.70, AP-related features were included in the optimization. Consequently, models that failed to generate at least one spike incurred a large penalty for missing features and were therefore excluded, biasing the population toward spiking responses at this stimulus intensity.

Another potential related limitation of our modelling is the too low variability of electrophysiological behaviour and the too high variability of ion channel parameters. Although our optimization allowed electrophysiological variability within two standard deviations from the mean, the resulting model population did not precisely match experimental variability. Comparison of mean e-feature values between experimental recordings (n = 29 neurons) and the model population showed excellent agreement (Pearson r = 0.996; S8 Figure left), confirming that mean electrophysiological behaviour is well reproduced. However, 170 out of 182 e-features showed SD ratios below 0.5, indicating that the variability across the model population is substantially lower than observed experimentally (S8 Figure right). This outcome is expected, given that optimizing models against mean feature targets with tight Z-score bounds tends to produce populations that cluster more tightly around the mean than the biological data do. This can be improved in the next optimizations of the available model population. In addition, the experimental variability of ion channel properties (expression, kinetics) in recorded early-birth CA1 pyramidal neurons is unknown and is likely lower than the relatively wide ion channel parameter variability allowed during our model optimization. A potential improvement in the form of a realistic reduction in the ion channel parameter variability might be achieved using Pareto optimization approaches that would also include energy-efficiency of conductance-based compartmental models [45].

Despite these limitations, the presented population of models provides a versatile resource for future studies. The availability of multiple validated parameter sets enables future systematic investigations of robustness, sensitivity, and energetic efficiency of intrinsic neuronal dynamics [36,45,54]. These models can be used for functional validation and hypothesis-driven predictions. Furthermore, they can be readily embedded into network simulations to study how intrinsic variability interacts with synaptic and circuit-level mechanisms. By explicitly capturing both parameter-level and feature-level diversity, this population-based framework enables analyses that are not accessible using a single representative model.

## Acknowledgements

We would like to thank Hermann Cuntz and Alexander Bird for useful discussions on morphological issues and for analysing data.

We thank the CINECA consortium (Bologna, Italy) and the HPC Devana system (Slovak Academy of Sciences, Bratislava, Slovakia) for providing access to their supercomputing resources.

AI was used to help with improvements to human-generated texts for readability and error-free grammar, spelling, punctuation and tone. The authors reviewed and verified the accuracy of the content.

## Data availability statement

All electrophysiological data used for optimization and the valid model parameters will be made available on the EBRAINS Knowledge Graph and the EBRAINS Live paper upon acceptance. All code used for model optimization, analysis, and figure generation will be archived on Zenodo and made available on GitHub upon acceptance.

## Supplementary Information

**S8 Figure:**
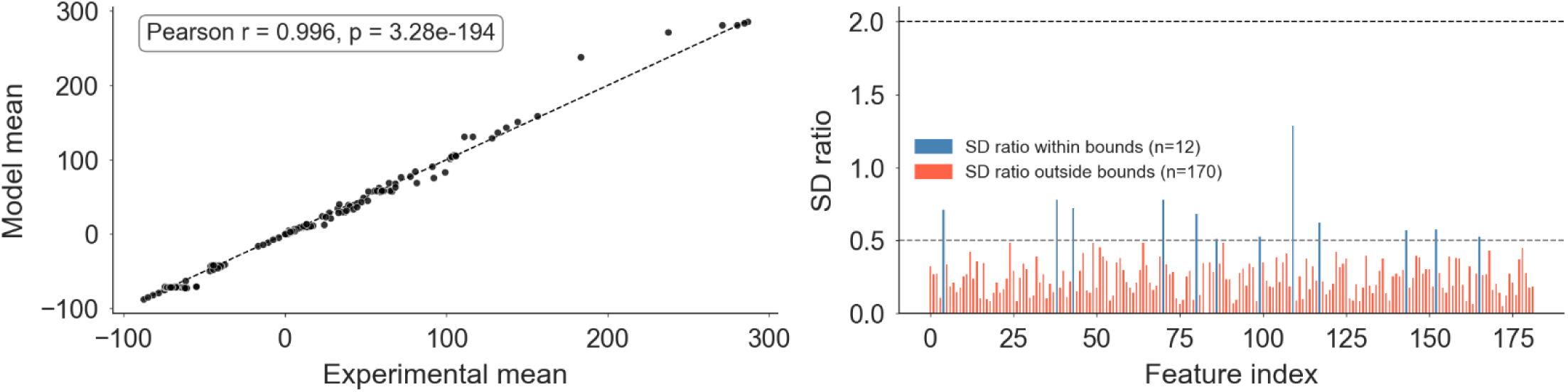
Comparison of electrophysiological feature distribution between experimental recordings and model populations. (Left) Scatter plot of mean feature values across all 182 e-features, comparing experimental recordings (n = 29 neurons) and model population (n = 3512 valid models). Each point represents one e-feature. The dashed diagonal line represents perfect agreement. Pearson r = 0.996, p = 3.28×10⁻¹⁹⁴. (Right) Standard deviation (SD) ratio (model SD / experimental SD) per feature across all 182 e-features. Dashed lines indicate the upper (2.0) and lower (0.5) bounds. Features outside these bounds (red) indicate substantial underestimation of experimental variability by the model population.

**S1 Table:** Values of the extracted e-features from experimental data for a 5-ms-long depolarizing current injection protocol.

| Feature / Input Current | 0.1 nA | 0.15 nA* | 0.55 nA | 0.5 nA* |
| --- | --- | --- | --- | --- |
| Maximum voltage from the voltage base | $7.61 \pm 1.40$<br>(29) | $11.37 \pm 2.17$<br>(29) | | |
| Decay time constant after stimulus | $10.61 \pm 11.54$<br>(29) | $8.35 \pm 4.96$<br>(29) | | |
| Voltage base | $-71.12 \pm 1.01$<br>(29) | $-71.22 \pm 1.29$<br>(29) | $-71.20 \pm 1.46$<br>(22) | $-71.20 \pm 1.14$<br>(22) |
| Steady state voltage | $-71.12 \pm 1.01$<br>(36) | $-71.22 \pm 1.29$<br>(29) | $-71.20 \pm 1.46$<br>(22) | $-71.20 \pm 1.14$<br>(22) |
| AP1 begin voltage | | | $-41.43 \pm 5.63$<br>(22) | $-40.41 \pm 6.57$<br>(19) |
| AP1 peak voltage | | | $65.20 \pm 6.23$<br>(22) | $66.00 \pm 6.47$<br>(22) |
| AP1 begin width | | | $3.07 \pm 0.85$<br>(22) | $3.32 \pm 0.79$<br>(19) |
| AP rise time | | | $0.63 \pm 0.06$<br>(22) | $0.64 \pm 0.07$<br>(19) |
| AP fall time | | | $2.64 \pm 0.41$<br>(22) | $2.65 \pm 0.47$<br>(22) |
| AP duration half width | | | $1.22 \pm 0.26$<br>(22) | $1.22 \pm 0.24$<br>(19) |
| AHP1 depth from peak | | | $110.79 \pm 24.80$<br>(32) | $116.11 \pm 6.88$<br>(22) |

\*Current step used in the generalization test

**S2 Table:** Values of the extracted e-features from experimental data for a 250-ms-long hyperpolarizing current injection protocol.

| Feature / Input Current | -0.025 nA | -0.05 nA* | -0.075 nA | -0.1 nA* | -0.125 nA |
| --- | --- | --- | --- | --- | --- |
| Voltage base | -71.12 ± 1.05<br>(29) | -71.48 ± 1.21<br>(29) | -71.75 ± 2.84<br>(29) | -71.95 ± 1.06<br>(29) | -71.87 ± 1.39<br>(29) |
| Steady state voltage | -71.12 ± 1.05<br>(29) | -71.48 ± 1.21<br>(29) | -71.75 ± 2.84<br>(29) | -71.95 ± 1.06<br>(29) | -71.87 ± 1.39<br>(29) |
| Steady state voltage stim end | -75.03 ± 1.56<br>(29) | -78.67 ± 2.28<br>(29) | -82.01 ± 3.02<br>(29) | -85.12 ± 3.59<br>(29) | -87.89 ± 4.12<br>(29) |
| Voltage deflection | -4.00 ± 1.36<br>(29) | -7.37 ± 2.24<br>(29) | -10.57 ± 2.84<br>(29) | -13.63 ± 3.61<br>(29) | -16.60 ± 4.20<br>(29) |
| Ohmic input resistance | 156.23 ± 55.03<br>(29) | 143.83 ± 45.40<br>(29) | 136.71 ± 36.96<br>(29) | 131.66 ± 35.84<br>(29) | 128.20 ± 32.42<br>(29) |
| Sag amplitude | 1.59 ± 0.66<br>(29) | 2.58 ± 1.47<br>(29) | 3.51 ± 1.91<br>(29) | 4.38 ± 2.35<br>(29) | 5.48 ± 2.92<br>(29) |
| Time constant | 24.11 ± 20.42<br>(29) | 16.97 ± 4.43<br>(29) | 15.81 ± 4.91<br>(29) | 14.46 ± 3.47<br>(29) | 13.20 ± 3.27<br>(29) |

\*Current step used in the generalization test

**S3 Table:**
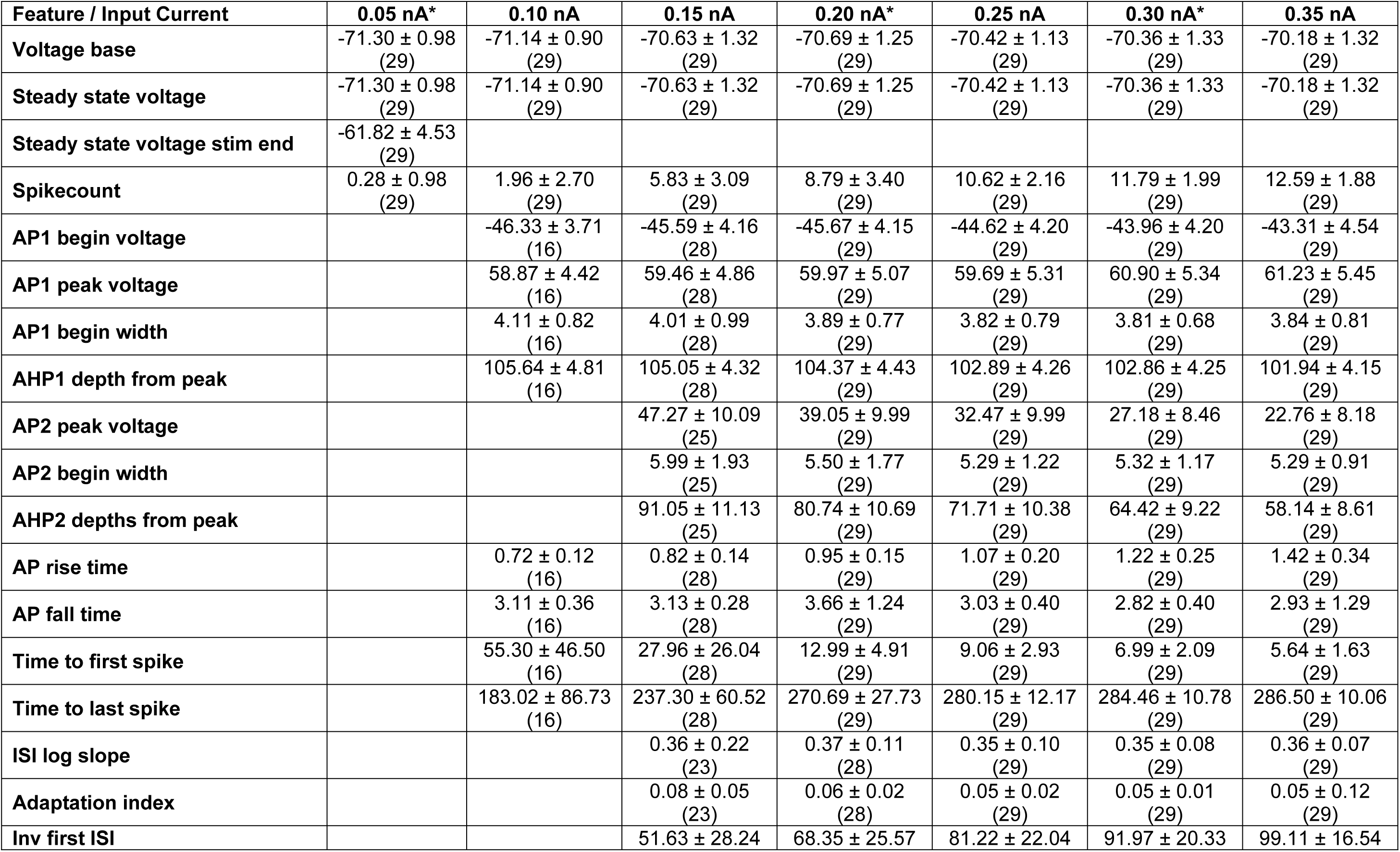
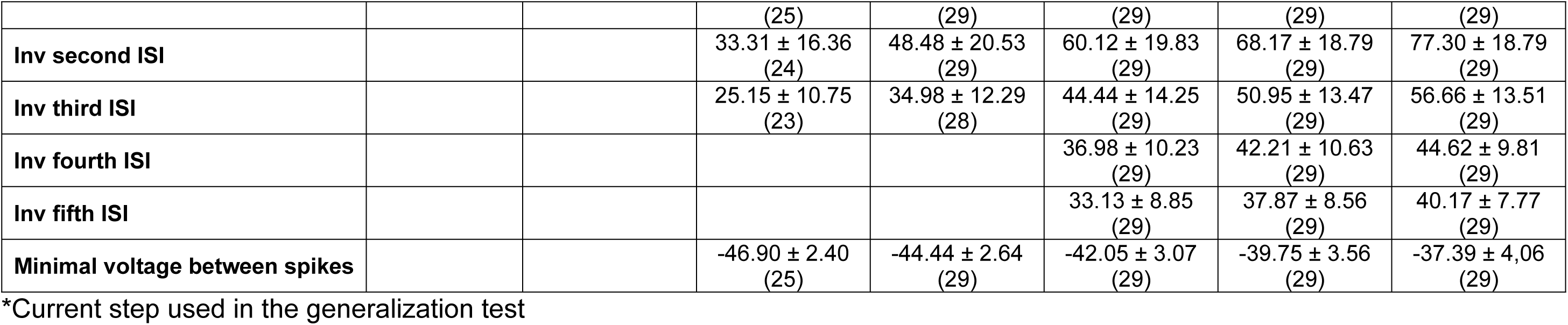
Values of the extracted e-features from experimental data for a 300-ms-long depolarizing current injection protocol.

**S4 Table:**
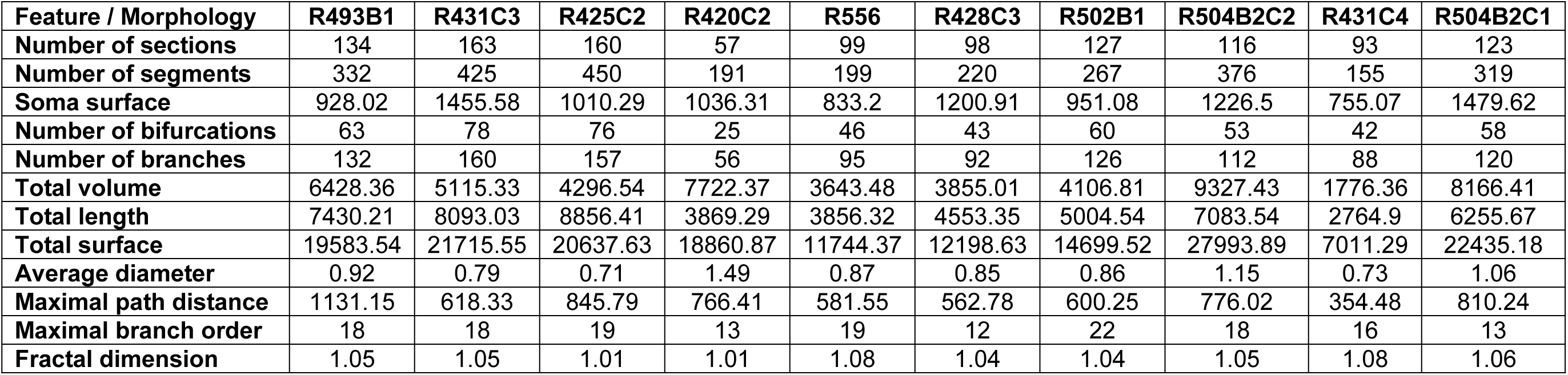
Morphological characteristics.

**S5 Table:** Constant parameters.

| Property | Value |
| --- | --- |
| Resting Membrane Potential (RMP, v_init) | -71.5 mV |
| Temperature (celsius) | 34 °C |
| Na<sup>+</sup> Nernst Potential (ena) | 65 mV |
| K<sup>+</sup> Nernst Potential (ek) | -90 mV |

**S6 Table:** Model parameters with their ranges used for optimization.

| Parameter (unit) | Section | Minimal value | Maximal value |
| --- | --- | --- | --- |
| *Passive properties* |  |  |  |
| Specific membrane capacitance (1μF/cm<sup>2</sup>) | All | 0.5 | 1.0 |
| Axial resistance (Ωcm) | Axon | 50 | 150 |
|  | Soma | 100 | 400 |
|  | Apical dendrites | 100 | 450 |
|  | Basal dendrites | 100 | 450 |
| Passive channel equilibrium potential (mV) | Axon | -80 | -60 |
|  | Soma, Dendrites | -75 | -60 |
| Passive channel maximal conductance (mS/cm<sup>2</sup>) | Axon | 0.00001 | 0.0004 |
|  | Soma | 0.0000015 | 0.0002 |
|  | Apical dendrites | 0.0000005 | 0.00005 |
|  | Basal dendrites | 0.00000025 | 0.0005 |
| *Active properties* |  |  |  |
| Transient sodium channel maximal conductance (mS/cm<sup>2</sup>) | Axon | 0.001 | 0.8 |
|  | Soma | 0.0025 | 0.2 |
|  | Apical dendrite | 0.00015 | 0.015 |
|  | Basal dendrites | 0.0075 | 0.2 |
| Half-activation voltage for the transient sodium channel, $\theta_a$ (mV) | All | -25 | -15 |
| Half-inactivation voltage for the transient sodium channel, $\theta_{i1}$ (mV) | All | -56 | -40 |
| Half-inactivation voltage for the transient sodium channel, $\theta_{i2}$ (mV) | All | -62 | -40 |
| Persistent sodium channel maximal conductance (mS/cm<sup>2</sup>) | Axon | 0.0001 | 0.0027 |
|  | Soma, Dendrites | 0.0000001 | 0.00002 |
| Half-activation voltage for the persistent sodium channel, $\nu_{half}$ (mV) | All | -64 | -50 |
| Delayed rectifier potassium channel maximal conductance (mS/cm<sup>2</sup>) | Axon | 0.0025 | 0.08 |
|  | Soma, Dendrites | 0.00005 | 0.0025 |
| Half-activation voltage for the delayed rectifier potassium channel, $v_{half}$ (mV) | All | 8 | 22 |
| M-type potassium channel maximal conductance (mS/cm<sup>2</sup>) | Axon | 0.000025 | 0.0175 |
|  | Soma | 0.000005 | 0.0012 |
| Half-activation voltage for the M-type potassium channel, $v_{half1}$ (mV) | Axon, Soma | -45 | -32 |
| Half-inactivation voltage for the M-type potassium channel, $v_{half2}$ (mV) | Axon, Soma | -47 | -37 |
| A-type potassium channel maximal conductance (mS/cm<sup>2</sup>) | Axon | 0.00025 | 0.3 |
| A-type potassium channel maximal conductance (mS/cm<sup>2</sup>) | Soma | 0.0005 | 0.05 |
| A-type potassium channel maximal conductance (mS/cm<sup>2</sup>) | Dendrites | 0.00015 | 0.02 |
| Half-activation voltage for the A-type potassium channel, $v_{halfn}$ (mV) | Axon, Soma | 4 | 16 |
|  | Dendrites | -1 | 6 |
| Half-inactivation voltage for the A-type potassium channel, $v_{halfn}$ (mV) | Axon, Soma | -63 | -51 |
| HCN channel maximal conductance (mS/cm<sup>2</sup>) | Soma, Dendrites | 0.000001 | 0.0001 |
| Half-activation voltage for the HCN channel, $v_{half1}$ (mV) | Soma, Dendrites | -120 | -95 |
| Slope of the HCN channel maximal conductance distribution | Soma, Dendrites | 0 | 0.03 |
| L-type calcium channel maximal conductance (mS/cm<sup>2</sup>) | Soma, Dendrites | 0.00000001 | 0.00002 |
| Half-activation voltage for the L-type calcium channel, $v_{halfm}$ (mV) | Soma, Dendrites | -1 | 5 |
| T-type calcium channel maximal conductance (mS/cm<sup>2</sup>) | Soma, Dendrites | 0.00000001 | 0.00002 |
| Half-activation voltage for the T-type calcium channel, $v_{halfm}$ (mV) | Soma, Dendrites | -33 | -23 |
| Half-inactivation voltage for the T-type calcium channel, $v_{halfh}$ (mV) | Soma, Dendrites | -80 | -70 |
| N-type calcium channel maximal conductance (mS/cm<sup>2</sup>) | Soma, Dendrites | 0.0000000025 | 0.000004 |
| Half-activation voltage for the N-type calcium channel, $v_{halfm}$ (mV) | Soma, Dendrites | -19 | -9 |
| Slow calcium-dependent potassium channel maximal conductance (mS/cm<sup>2</sup>) | Soma,<br>Dendrites | 0.00001 | 0.0004 |
| Calcium-activated potassium channel maximal conductance (mS/cm<sup>2</sup>) | Soma,<br>Dendrites | 0.000001 | 0.0004 |

**S7 Table:** Pairwise comparisons of total scores between morphologies. P-values are reported for all morphology pairs following Dunn’s post hoc test with Holm correction, applied after a significant Kruskal-Wallis test (H=1251.970, p=7.293×10^-264^). Green shading indicates non-significant differences (p > 0.05); all other pairs differ significantly.

| Morphology | R493B1 | R431C3 | R425C2 | R420C2 | R556 | R428C3 | R502B1 | R504B2C2 | R431C4 | R504B2C1 |
| --- | --- | --- | --- | --- | --- | --- | --- | --- | --- | --- |
| R493B1 | 1.0 | $4.666 \times 10^{-37}$ | $7.65 \times 10^{-53}$ | $3.032 \times 10^{-26}$ | $7.286 \times 10^{-03}$ | $4.699 \times 10^{-09}$ | $5.330 \times 10^{-08}$ | $3.194 \times 10^{-29}$ | $1.314 \times 10^{-10}$ | $7.666 \times 10^{-34}$ |
| R431C3 | $4.666 \times 10^{-37}$ | 1.0 | $5.130 \times 10^{-03}$ | $3.361 \times 10^{-01}$ | $3.002 \times 10^{-14}$ | $1.719 \times 10^{-54}$ | $9.125 \times 10^{-51}$ | $3.964 \times 10^{-90}$ | $2.303 \times 10^{-02}$ | $6.875 \times 10^{-89}$ |
| R425C2 | $7.65 \times 10^{-53}$ | $5.130 \times 10^{-03}$ | 1.0 | $4.357 \times 10^{-04}$ | $9.126 \times 10^{-23}$ | $4.210 \times 10^{-69}$ | $9.621 \times 10^{-65}$ | $5.349 \times 10^{-107}$ | $2.360 \times 10^{-05}$ | $6.821 \times 10^{-104}$ |
| R420C2 | $3.032 \times 10^{-26}$ | $3.361 \times 10^{-01}$ | $4.357 \times 10^{-04}$ | 1.0 | $2.603 \times 10^{-10}$ | $7.798 \times 10^{-44}$ | $6.724 \times 10^{-41}$ | $1.133 \times 10^{-75}$ | $1.411 \times 10^{-01}$ | $6.930 \times 10^{-77}$ |
| R556 | $7.286 \times 10^{-03}$ | $3.002 \times 10^{-14}$ | $9.126 \times 10^{-23}$ | $2.603 \times 10^{-10}$ | 1.0 | $3.395 \times 10^{-13}$ | $4.865 \times 10^{-12}$ | $5.540 \times 10^{-33}$ | $1.917 \times 10^{-04}$ | $9.640 \times 10^{-38}$ |
| R428C3 | $4.699 \times 10^{-09}$ | $1.719 \times 10^{-54}$ | $4.210 \times 10^{-69}$ | $7.798 \times 10^{-44}$ | $3.395 \times 10^{-13}$ | 1.0 | $7.867 \times 10^{-01}$ | $6.872 \times 10^{-07}$ | $1.120 \times 10^{-23}$ | $7.504 \times 10^{-11}$ |
| R502B1 | $5.330 \times 10^{-08}$ | $9.125 \times 10^{-51}$ | $9.621 \times 10^{-65}$ | $6.724 \times 10^{-41}$ | $4.865 \times 10^{-12}$ | $7.867 \times 10^{-01}$ | 1.0 | $2.348 \times 10^{-07}$ | $3.340 \times 10^{-22}$ | $2.279 \times 10^{-11}$ |
| R504B2C2 | $3.194 \times 10^{-29}$ | $3.964 \times 10^{-90}$ | $5.349 \times 10^{-107}$ | $1.133 \times 10^{-75}$ | $5.540 \times 10^{-33}$ | $6.872 \times 10^{-07}$ | $2.348 \times 10^{-07}$ | 1.0 | $3.006 \times 10^{-45}$ | $5.337 \times 10^{-02}$ |
| R431C4 | $1.314 \times 10^{-10}$ | $2.303 \times 10^{-02}$ | $2.360 \times 10^{-05}$ | $1.411 \times 10^{-01}$ | $1.917 \times 10^{-04}$ | $1.120 \times 10^{-23}$ | $3.340 \times 10^{-22}$ | $3.006 \times 10^{-45}$ | 1.0 | $7.103 \times 10^{-50}$ |
| R504B2C1 | $7.666 \times 10^{-34}$ | $6.875 \times 10^{-89}$ | $6.821 \times 10^{-104}$ | $6.930 \times 10^{-77}$ | $9.640 \times 10^{-38}$ | $7.504 \times 10^{-11}$ | $2.279 \times 10^{-11}$ | $5.337 \times 10^{-02}$ | $7.103 \times 10^{-50}$ | 1.0 |

